# No Evidence for Stronger Brain–Behavior Associations in the Bimanual Motor Network of Older Adults

**DOI:** 10.1101/2025.11.15.688248

**Authors:** Mikael Novén, Jesper Lundbye-Jensen, Henrik Lundell, Hartwig Roman Siebner, Anke Ninija Karabanov

## Abstract

Bimanual motor performance declines with age, is accompanied by cortical thinning and alterations in white matter microstructure within motor control networks. Aging is also associated with increased interindividual variability in behavioural and structural markers, and some studies report stronger brain–behaviour associations in older compared to younger adults. However, evidence for this remains mixed, and it is unclear whether young adults express similar relationships under comparable task demands. We tested whether bimanual performance is related differently to structural brain properties in younger and older adults. Twenty-three younger (22–27 years) and twenty-three older adults (65–70 years) performed a visually guided bimanual pinch-force task and underwent whole-brain structural magnetic resonance imaging (MRI). Fractional anisotropy (FA) and cortical thickness (CTh) were extracted from a pre-defined bilateral visuo-motor network and compared between groups. In regions showing significant age-related differences, we tested whether and how FA and CTh values correlates with bimanual performance within each age group. Older adults showed lower FA in the anterior portions of the bilateral superior longitudinal fasciculus (SLF) III and right inferior fronto-occipital fasciculus (IFOF), and lower CTh across most visuomotor regions, along with lower bimanual performance compared to young adults. Structure–function analyses in these areas revealed that bimanual performance correlated positively with FA in the anterior segment of the right SLF III in younger but not older adults. Furthermore, larger hemispheric asymmetries in cortical thickness between the dominant and non-dominant SMA correlated positively with larger intermanual performance differences between the dominant and non-dominant hand in younger but not older adults. These findings suggest that intrahemispheric white-matter integrity and interhemispheric cortical balance support efficient bimanual control in early adulthood, but they provide no evidence that structure–function relationships increase with age.

## Introduction

The ability to coordinate both hands is essential for many activities of daily living. However, bimanual coordination deteriorates across the adult lifespan (Boisgontier et al., 2018). Older adults show longer movement times, higher performance variability, and reduced movement accuracy (Kang & Cauraugh, 2022; Krehbiel et al., 2017; Roman-Liu & Tokarski, 2021; Rudisch et al., 2020; Vieluf et al., 2015). This decline in bimanual function is accompanied by well-documented age-related changes in cortical thickness in key nodes of the bimanual visuomotor network, including the primary motor cortex (M1), premotor cortex, supplementary motor cortex (SMA) and posterior parietal cortex (Lemaitre et al., 2012), as well as degeneration of white matter microstructure of motor-related white matter tracts such as the corticospinal tract (CST), the corpus callosum (CC), or the SLF (Oschwald et al., 2021; Sexton et al., 2014).

Studies have shown that better bimanual performance is associated with higher callosal white matter integrity and greater grey matter volume in the cerebellum and sensorimotor areas across age groups (Cobia et al., 2024; Johansen-Berg et al., 2007; Koppelmans et al., 2015; Serbruyns et al., 2015). However, less is known about whether aging affects the relationship between brain structure and motor performance. Longitudinal aging studies suggest that age-related atrophy of grey and white matter is linked to declines in manual function and demonstrate that better manual dexterity in older adults is associated with less atrophy of both grey and white matter across several cortical and subcortical regions (Dougherty et al., 2024, Zivari Adab et al., 2020)). Some cross-sectional studies have directly compared the relationship between brain structure and motor performance between older and younger adults. Some suggest age-related differences in the relationship between callosal white matter integrity and bimanual performance measures (Serbruyns et al., 2015; Fujiyama et al., 2016) and between grey matter volume in sensori-motor areas and the cerebellum and motor performance (Boisgontier et al., 2018), with higher brain-performance correlations in older adults. However, this literature is not uncontested; other studies have not been able to demonstrate that age is a mediating factor in brain-behavior relationships (Ruitenbeek, 2017), and others found stronger correlations between motor performance measures, such as reaction time and callosal white matter integrity in younger compared to older adults (Madden et al., 2004).

A limited number of studies have included direct comparisons in structure–motor performance associations between different age groups. Data by Boisgontier and coworkers compared performance in a bimanual visuomotor task across different age groups. They found that bimanual coordination performance was predicted by grey matter volume in the cerebellum in the groups who showed the worst task performance: children and adolescents, and adults over 60, with primary sensorimotor cortex additionally contributing in the older group. In contrast, young adults, who showed the least individual variability, had no significant structure–performance correlations (Boisgontier et al., 2018). Another study found stronger and more extensive correlations between various aspects of bimanual coordination and sub-regions of the corpus callosum in older compared to younger adults (Serbruyns et al., 2015).

Among diffusion tensor metrics, FA is considered a sensitive marker of age-related white-matter decline, showing robust tractwise effects (Behler et al., 2021; Matijevic & Ryan, 2021) and has therefore often been chosen as the main outcome variable. Previous studies on the associations between white FA and bimanual control have almost exclusively focused on transcallosal tracts, which may mean that other important tracts are overlooked. Functional MRI studies suggest that dynamic, visually guided bimanual movements rely heavily on froto-parietal connections like the SLF (Karabanov et al., 2023). Older adults show performance decline in these tasks, but the age-related differences in brain connectivity between age groups in dynamic, visually guided bimanual movements remain largely unexamined (Wulff-Abramsson et al., 2025; Zvornik et al., 2024).

Task differences may also influence age-dependent differences in the structure-performance relationship. Age-related changes in the structure–performance relationships vary considerably by task type and demands (Razlighi et al., 2017). For example, a large study (>600 participants) found that the correlations between memory and white matter integrity in hippocampal pathways declined after 60 years, while correlations with language or fluid intelligence remained stable across the lifespan. This highlights that for some tasks, structure–performance coupling also weakens in later life (de Mooij et al., 2018). These findings suggest that age may also weaken structure–performance coupling in certain domains, potentially due to dedifferentiation processes that reduce the regional specificity of brain function.

This study aims to investigate the relationship between brain structure, bimanual performance, and ageing using an approach that does not a priori focus on the corpus callosum. Instead, the study first identifies which structures show age-related differences in white matter microstructure (FA) and cortical thickness within the brain’s bimanual control network. Second, it examines, within these regions, whether the association between structural measures and bimanual performance shows different patterns depending on the age group. Based on prior evidence of increased correlations between bimanual performance markers and callosal integrity, we expect stronger structure–performance associations in older adults.

## Methods

### Participants

Twenty-nine older (OA; 65-70 y.o.) and 25 younger (YO; 22-27 y.o.) healthy adults with no history of neuropsychiatric disease and not taking any neuroactive medication were recruited to participate in this study. All included participants had normal cognitive function assessed by the Montreal Cognitive Assessment (MOCA) (Nasreddine et al., 2005) and were classified as right-handed, as assessed by the Edinburgh Handedness Inventory (EHI) (Oldfield, 1971). Three OA were excluded due to MOCA scores below our exclusion criteria of 25, 1 OA due to claustrophobia in the MRI scanner, 2 OA due to incidental findings, and 2 YA due to faulty scans. The final sample was thus 23 subjects in each age group. All participants gave written informed consent before participation and were reimbursed for their participation. The study was approved by the Regional Committee on Health Research Ethics of the Capital Region in Denmark (De Videnskabsetiske Komitéer-Region H; Journal-nr: H-21025010).

### Experimental procedure

The experiment was split between two days of testing. On day one, participants were informed about the details of the study, gave their informed consent to participate, and completed a number of screening tests. On the second day, participants were welcomed to the scanning facility, underwent MRI safety screening, were instructed on the bimanual task and scanning procedure, before finally undergoing MRI scanning.

After signing consent, all participants performed a selection of motor-cognitive background tests to verify that cognitive and motor functioning fell within an expected functional range and to identify potential group differences in mental and physical functioning. Participants were screened for cognitive functioning (MOCA) and right-handedness (EHI). Grip- and pinch strength, measured by hydraulic gauges (Baseline® Lite Hydraulic hand dynamometer Prod. Nr: 12-0241 and Baseline® Lite Hydraulic Pinch Gauge Prod. Nr: 12-0226, Elmsford, NY, U.S.A.), was used to aid calibration of the pinch force task and to quantify upper limb strength. Participants were also screened for their general physical fitness to ensure homogeneous physical activity across age groups. Aerobic fitness was assessed based on VO2-max estimations from seismocardiographic recordings using Seismofit® (VentriJect 2022, 2900 Hellerup, Denmark). Subjects also completed a 30-second sit-to-stand test (Lein Jr et al., 2022) and completed the International Physical Activity Questionnaire (Craig et al., 2003). Bimanual dexterity was assessed using the Purdue Pegboard test (Tiffin & Asher, 1948). Finally, a subset of the Cambridge Neuropsychological Testing Automated Battery was administered, specifically the paired associates learning, rapid visual information processing with three targets, and reaction time in five-choice mode tasks (Fray et al., 1996). The most relevant characteristics from day 1 are summarized in Table 1, while other measures are reported elsewhere (Weakly et al, in submission). VO2 max estimates were within normal values for the age groups in Denmark (20-29 y.o. Male: 54.4 ± 8.4, Female: 43.0 ± 7.7; 60-69 y.o. Male: 39.2 ± 6.7, Female: 31.1 ± 5.1) (Loe et al., 2013).

**Table 1.** Demographic and background measurements for included participants. If nothing else is indicated, reported values are group averages ± standard deviation. The groups did differ significantly in VO2 max (t(42.8)=6.42, p=9.10e-08), MOCA score (t(40.0)=2.76, p= 0.00879), and pegboard assemblies (t(43.9)=5.27, p=3.99e-06). The asterisk to the right of each measure denotes significant group differences.

|  | Young group<br>(22-27 y.o.) | Older group (65-70<br>y.o.) |
| --- | --- | --- |
| Sex (F/M) | 11/12 | 11/12 |
| Age in years (range)* | 24.7 (22 – 27) | 67.3 (65 – 70) |
| BMI | 23.6 ± 3.6 | 25.4 ± 4.4 |
| Pegboard Assemblies * | 30.0 ± 5.6 | 21.2 ± 5.9 |
| Max pinch force Left | 8.8 ± 2.4 | 8.0 ± 2.1 |
| Max pinch force Right | 9.2 ± 2.2 | 8.4 ± 2.6 |
| 30s Sit-to-stand | 17.8 ± 3.6 | 17.2 ± 5.4 |
| VO2 max estimate *<br>/mL/kg/min | 46.9 ± 8.4<br>(average-high<br>for age group) | 32.1 ± 7.1<br>(average-high for age<br>group) |
| MOCA score * | 29.0 ± 0.9 | 28.1 ± 1.3 |

### Bimanual pinch grasp task

After the participants underwent MRI safety screening on day 2, participants were placed in the scanner where they were introduced to the pimanual pinch task which consisted of dynamic transitions between diffenrnet pinch force levels requiring bimanual coordination (asymmetric and symmetric blocks) and blocks of static hold at a delicate force level, requiring stable fine force control (Figure 1; see also (Wulff-Abramsson et al., 2025)). When lying in the MRI scanner, participants had one force sensor, 3 cm in diameter, in a thumb pinch position in each hand (Figure 1A). Pinch force controlled the diameter of yellow semicircles on the corresponding half of a screen visible through a mirror attached to the head coil. Sensor data was sampled at 1024 Hz, but the displayed force trace was smoothed, i.e., averaged over the last 100 samples, to minimize electrical noise. Before the task started, the subject’s max voluntary contraction (MVC) was measured for each hand, and the hand-averaged MVC was used as a reference value for the target force levels of both hands. The session included four runs built up by a static baseline hold condition at 5% MVC for 20 seconds, followed by 6 symmetric and 6 asymmetric condition blocks in an interleaved order before the run was ended by another baseline hold (Figure 1C). Each block consisted of 20 trials á 2 seconds (± 100ms jitter) that directly followed each other, in which the target force for each hand was either 2.5, 5, 7.5, or 10% of the participant’s MVC, indicated by a blue target semicircle (figure 1B). Each block was separated by a three-second pause. Participants were allowed to rest between each run. Participants performed the task during four functional echo planar imaging sequences used for fMRI. While the fMRI data is reported elsewhere (Weakly et al, submitted), we use the behavioral data acquired during these scans for performance correlations with structural markers.

**Figure 1.**
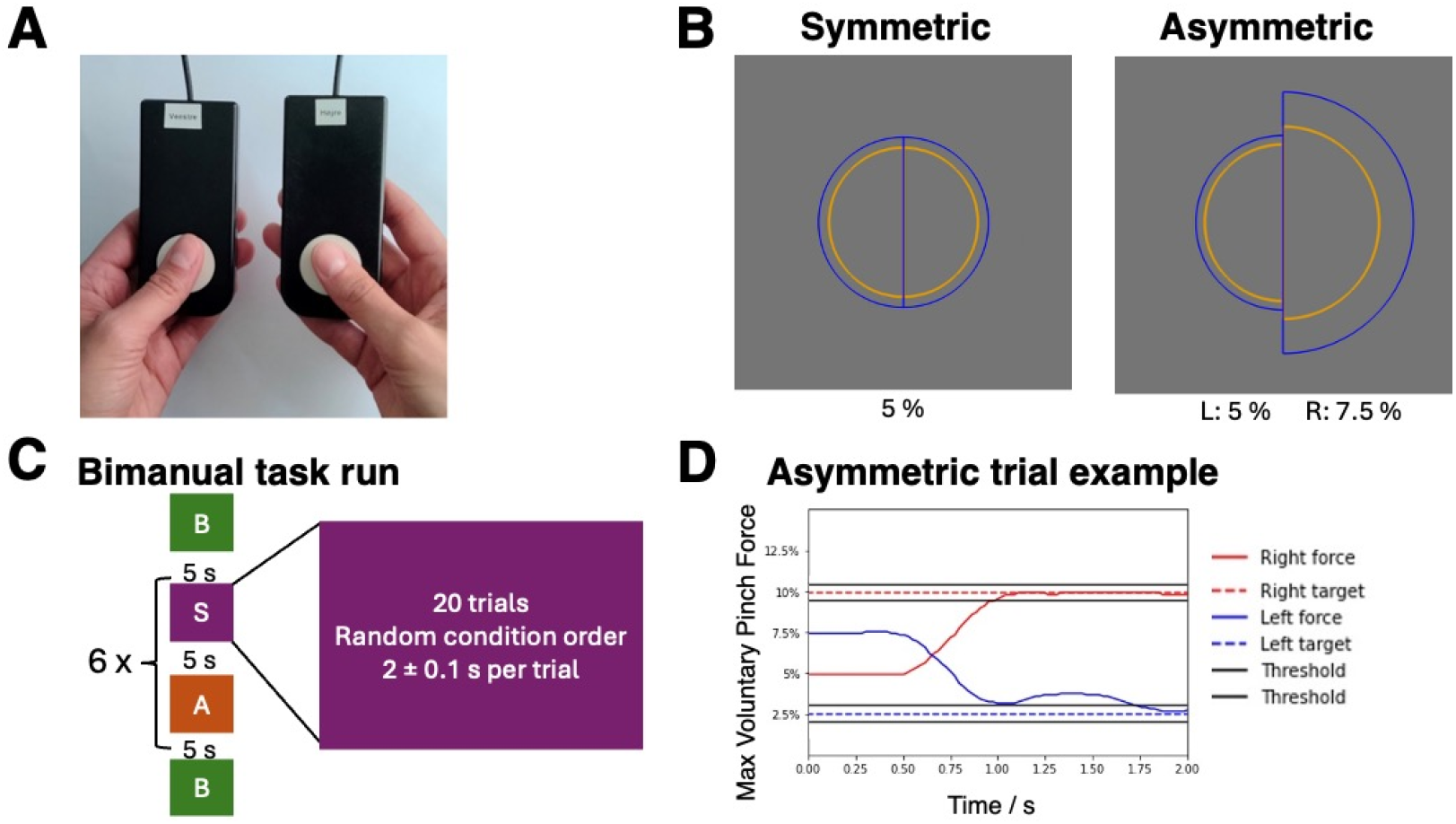
Task overview. A: Handheld pinch force sensors are used for maximum voluntary contraction measurement and the pinch force task. B: Example symmetric and (mirror) asymmetric trial types used in the task blocks. Pinch force applied from the left and right hands controlled the radius of the corresponding semicircle (yellow). Percentages are the target force levels (blue), relative to each subject’s maximum voluntary contraction. C: Overview of one experimental run. D: Example performance from an asymmetric trial. Red and blue lines correspond to the left and right hands, respectively. Solid black lines indicate the accuracy threshold for the Time-on-Target measure.

### Extraction of behavioral measures

First, the force data were log-transformed to comply with Weber’s law of perception of change (Fechner, 1860). To effectively capture bimanual performance, the time on target (ToT) was calculated as the percentage of trial time spent within 0.5% MVC of the target line (Figure 1D). ToT is sensitive to both speed and accuracy and is thus the primary measure of interest for dynamic task conditions that require subjects to reach the target as quickly as possible and maintain a stable force once the target is reached. We decided to calculate TOT as a unified behavior variable across symmetric and asymmetric blocks because previous work from our group has shown that individual performance within symmetric and asymmetric blocks is highly correlated (Weakly et al., in prep.). We also decided to average TOT across both hands, as it correlates highly between left and right hands (see Supplementary Figure 1). However, as we were also interested to gain insight in bimanual coordination we additionaly quantify the intermanual synchronization, we also calculated the absolute differences between the right and left hand for ToT, as also done in our previous studies (Karabanov et al., 2023).

### Magnetic Resonance Imaging

Data acquisition was performed at the Danish Research Centre for Magnetic Resonance, Hvidovre Hospital, Denmark, using a 3T MR scanner (Philips Achieva, Best, Netherlands) with a 32-channel array receive coil. While the primary outcome measures in this study were cortical thickness and white matter microstructure, the scanning protocol also included four EPI sequences for BOLD-fMRI. For completeness, the full acquisition protocol is described here. The scanning session started with the acquisition of a T1-weighted image (MPRAGE; FOV: 245 mm; 245 sagittal slices, TR/TE: 6.0 / 2.7 ms; resolution 0.85×0.85×0.85mm^3^; flip angle: 8°; TI: 755.9 ms) used for coregistration and quantification of cortical thickness. Four repetitions of a T2*-weighted echo planar imaging (EPI) sequence utilizing gradient echo (FOV: 190 mm; 42 transverse slices acquired in interleaved order, TR/TE: 2000/30 ms; in-plane resolution:3×3 mm^2^, 3mm slices, no slice gap, flip angle: 90°), interleaved with four identical EPI sequences with reversed phase encoding for susceptibility distortion correction, used for functional MRI analysis (reported elsewhere: Weakly, submitted). A T2-weighted image (TSE; FoV: 245×245 mm^2^; 245 sagittal slices, TR/TE: 2.5 s / 270 ms; resolution 0.85×0.85×0.85mm^3^; flip angle: 90°.; TI: 755.9 ms), a Fluid-Attenuated Inversion Recovery image (FLAIR; 256 mm; 202 sagittal slices, TR/TE: 4.8 s / 330 ms; resolution 1×1×1mm^3^; flip angle: 90 deg.), used for pathology assessments. Finally, one conventional diffusion-weighted imaging (DWI) sequence (spin-echo EPI, axial FOV: 224×224 mm; 66 slices; TR/TE: 7970 /90; resolution: 2×2×2mm^3^, flip angle: 90°, b = 0 and 2000 s/mm^2^ with 48 gradient directions) with additional 5 b = 0 s/mm^2^ volumes acquired using reversed phase encoding for susceptibility error correction, was acquired and used for measuring white matter microstructure parameters. One older individual could not complete the diffusion-weighted sequence.

### Bimanual motor control network segmentation

To define the task-relevant cortical areas and white matter tracts, we used data delineating the functional network that is activated during this specific task (Karabanov et al., 2023). The resulting bimanual reaching network was composed of the bilateral primary motor cortex hand area (M1hand), bilateral dorsal premotor cortex (PMd), bilateral supplementary motor area (SMA), and bilateral visual area 5, with coordinates taken from our previous work (Karabanov et al., 2023). The cortical regions of interest (ROIs) used in his study were chosen as the closest counterparts to these coordinates in Glasser’s multimodal atlas (Glasser et al., 2016), except for the M1hand ROI, which was taken from another publication in order to focus on the hand-knob instead of the entire M1 (Dubbioso et al., 2021). To further define the task-relevant tracts, the TractSeg (see method below) output tracts that combined two or more of the cortical network nodes were used as tracts of interest (TOIs) for subsequent analyses. The identified TOIs were the fourth corpus callosal tract (CC_4), the bilateral superior longitudinal fasciculus (SLF) parts I, II, and III, and the inferior fronto-opercular fasciculus (IFOF). The CC_4 was further refined by using bilateral M1hand ROIs from (Dubbioso et al., 2021) as inclusion masks.

### White matter microsctruture – preprocessing and analysis

The DWI data were used to segment TOIs and to quantify their microstructure. DWI data was preprocessed by registering the reversed phase encoding data to the main DWI b0 image using FSL’s FLIRT tool (Jenkinson et al., 2002), denoising all DWI data using *dwidenoise* in MRTrix3 (Cordero-Grande et al., 2019; Veraart et al., 2016), brain extraction using FSL’s BET (Smith, 2002), and finally susceptibility and eddy current distortion correction using FSL’s TOPUP (J. L. R. Andersson et al., 2003) and eddy (J. L. Andersson & Sotiropoulos, 2016). Next, we used FSL’s *dtifit* to obtain whole-brain fractional anisotropy (FA) and mean diffusivity (MD) maps. The maps capture different aspects of the white-matter microstructure. FA provides a measure of axonal structure and alignment. Thereby, it is sensitive to axonal integrity and myelination (Curran et al., 2016) and it is our primary measure for white matter microsctructure. MD quantifies the average mobility of water molecules in the voxel along all directions of measurement. It is thus sensitive to the general restriction of water in the tissue. Although FA was our primary, we also analyzed MD to provide complementary information on overall tissue diffusivity. To obtain the TOIs, we used TractSeg, an openly available automatic segmentation tool based on a convolutional neural network trained on data from the Human Connectome Project (Wasserthal et al., 2018). This provided anatomically valid segmentations of the white matter trancts of interest. FA and MD were sampled along 100 segments along each TOI using TractSeg’s *tractometry* function (Wasserthal et al., 2020). As FA was the primary measure of microstructural integrity in this study all results related to MD can be found in the supplementary material.

### Cortical thickness - preprocessing and analysis

Cortical surface reconstruction and volumetric segmentation were performed using the FreeSurfer image analysis suite v7.2 (B. Fischl et al., 2001; Dale et al., 1999; Fischl et al., 2004; Fischl & Dale, 2000). Briefly, cortical thickness (CT) was calculated as the distance between the pial (i.e. the border of the cerebral cortex and the cerebrospinal fluid) and white (i.e. the border between the white and grey matter) surfaces. The resulting whole-brain CTh maps were registered to FreeSurfer common space *fsaverage* and used for group difference analyses. Moreover, the average CTh within each cortical ROI using the FreeSurfer tool *mri_segstats* was calculated for each participant for specific group difference and correlational analyses.

### Statistical analyses

#### Behavioral Performance

Group differences in ToT and bimanual symmetry were tested using independent t-tests in each of the task conditions with an alpha thereshold of p < 0.05. Shapiro-Wilk normality tests, t-tests and correlation analyses were performed with standard commands (*shapiro*.*test, t*.*test* and *cor*.*test*, respectively) in R version 4.1.2 (R Core Team, 2021). If normality was not confirmed by p > 0.05 in the Shapiro-Wilk test, Mann-Whitney test (*wilcox*.*test*) and Spearman correlation methods were used instead of t-tests and Pearson correlation tests, respectively, with standard commands in R.

#### Group Differences in White Matter Microstructure and performance-microstructure correlations

To identify the TOI sections that were impacted by ageing, each TOI was divided into 100 segments. Diffusion parameters (i.e. FA and MD) were compared between groups for each tract segment using t-tests and family-wise error correction for multiple comparisons along the tract (alpha threshold 0.05) (Wasserthal et al., 2020; Yeatman et al., 2012). In case of significant group differences that extended at least 5 segments along the tract, averages of the diffusion parameter from within the significant tract segment were extracted and correlated with TOT for the significant sections separately for each group. If normality was not confirmed by p > 0.05 in the Shapiro-Wilk test, Spearman correlation methods were used instead of Pearson. The p-values were Bonferroni corrected across the three segments that should significant group diffenreces.

#### Group Differences in Cortical Thickness and performance-CTh correlations

To identify the cortical areas that were impacted by ageing we compared CTh between groups across the whole brain using *mri_glmfit* in FreeSurfer, including permutation test for multiple comparison correction (10^4^ permutations, p threshold: 0.001, cluster-wise p threshold: 0.05) and threshold-free cluster enhancement (Winkler et al., 2014). Average CTh from the bimanual control network ROIs was Pearson correlations with TOT were calculated for the significant ROIs separately for each group in R. If normality was not confirmed by p > 0.05 in the Shapiro-Wilk test, Spearman correlation methods were used instead of Pearson. P values were Bonferroni corrected across the 7 ROIs that showed significant group differences.

#### Correlations between hand synchrony and hemispheric asymmetries

To explore whether there were any correlations and hand synchrony in performance and hemispheric assymetries in brain structure, the absolute difference in performance measures between hands was Pearson correlated in R with the absolute difference in brain structure measure between hemispheres. For cortical ROIs, this meant the absolute difference in average cortical thickness between the left and right hemispheric homologues. For the tracts of interest, it meant the absolute difference in average diffusion parameter between the left and right hemispheric homologues. The CC tract was split into a left and right hemispheric component and considered hemispheric homologues. If normality was not confirmed by p > 0.05 in the Shapiro-Wilk test, Spearman correlation methods were used instead of Pearson. The p-values were Bonferroni corrected across the 4 left-right ROIs where at least one hemisphere has shown significant group differences.

### Tests for differences in correlation

If performance and brain structure measures correlated for any of the groups, the “cocor” R package was used to test whether the correlations within each group was significantly different, based on the computation of confidence intervals (Diedenhofen & Musch, 2015). All correlation difference tests were two-tailed, the alpha threshold was set to 0.05 and the confidence level to 95%.

## Results

### Bimanual task performance

Performance during the dynamically changing task was measured by the fraction of time spent on target (ToT). Both ToT and absolute hand difference in ToT were normally distributed (Shapiro-Wilks test, p>0.05). Younger adults showed significantly higher ToT values compared to older adults (M_YA_=40.0%, M_OA_=29.5%, p=6.37e-11; figure 2A). Older adults showed a significantly larger absolute difference between the left and right hand (M_OA_=18.1%, M_YA_=16.8%, p=0.0333; figure 2B). ToT correlated with manual dexterity as measured by the Purdue Pegboard Assembley task when combining both groups (Spearman’s *rho* = *0*.*354, p<0*.*001*). However, this relationship was largely attributable to between-group differences. There was no significant correlation within either in the young (Spearman’s rho = -0.055, p=0.716) or older (Spearman’s rho = -0.085, p=0.575) age group.

**Figure 2.**
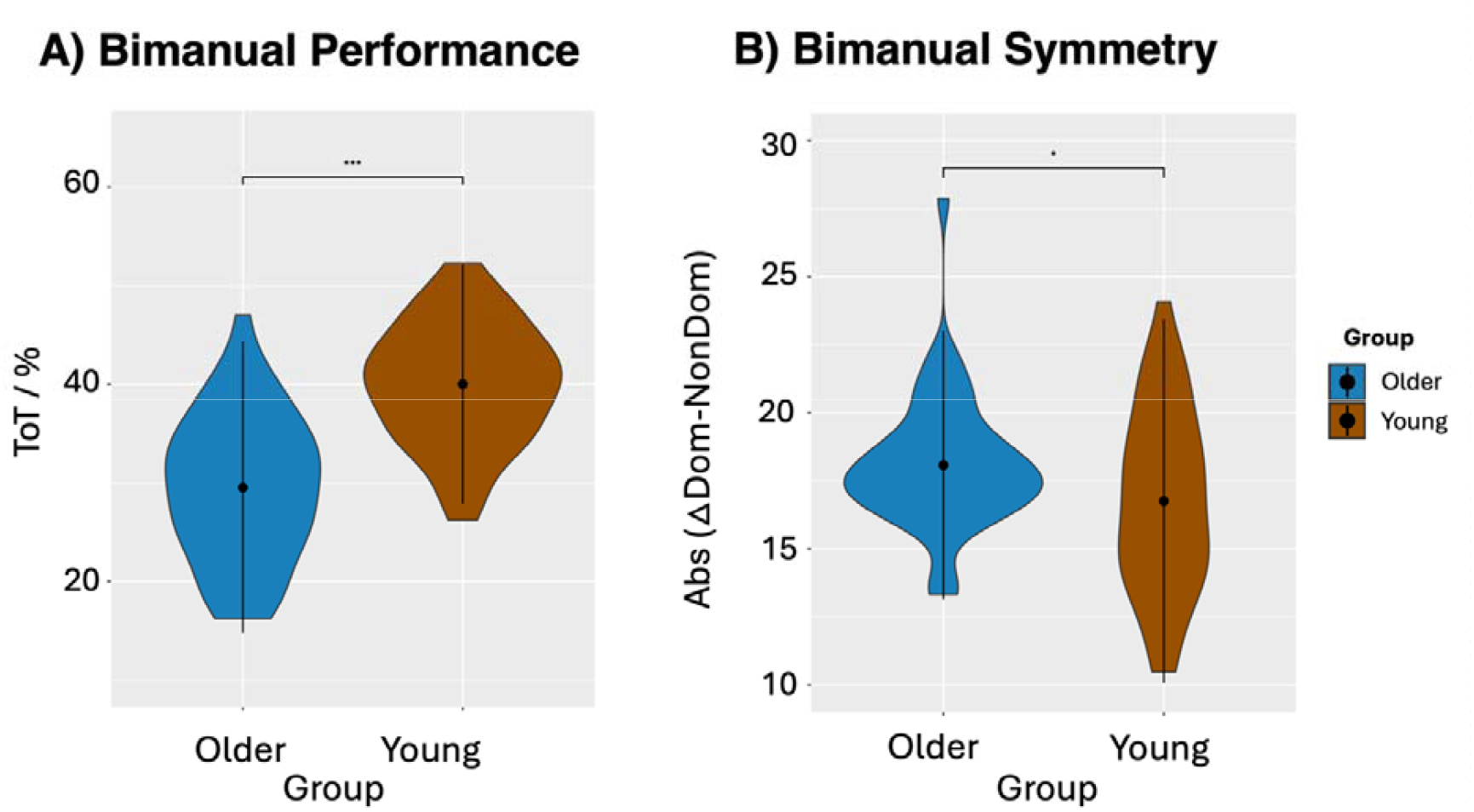
Time on Target in the bimanual motor task for older and younger adults. A) average time on target (ToT) of both hands during a trial as a measure of task performance B) Absolute difference in ToT in the dominant (Dom) and non-dominant (NonDom) hand as a measure of bimanual symmetry. Results for older adults are shown in blue, results for younger adults are shown in red. Stars denote significane values: * p<0.05, ** p<0.01, p<0.001***

### Group differences in Fractional Anisotropy and Cortical Thickness

The younger age group had greater FA in segments of the right IFOF, and the bilateral SLF III when compared with the older adults (figure 3). No other tracts showed a significant group difference. Mean diffusivity was significantly lower for younger compared to older adults in the bilateral IFOF and right SLF I (supplementary material, figure 3).

**Figure 3.**
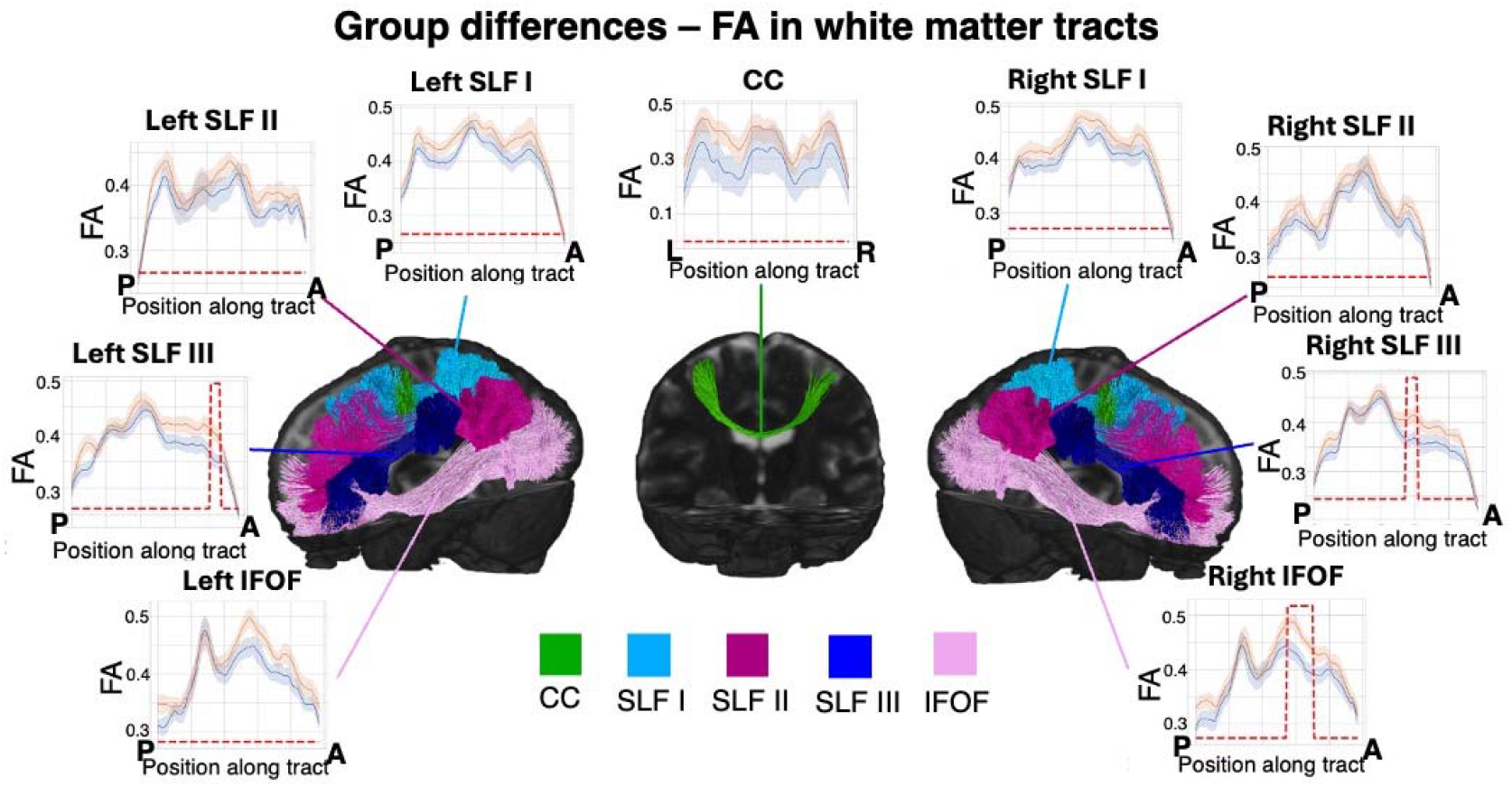
Group differences in Fractional Anisotropy (FA) in the tracts connecting the bimanual control nodes. FA values of older adults are displayed in blue while FA values of the younger adults are shown in red. Position labels on the x-axis indicate left (L), Right (R), Anterior (A), and Posterior (P). The dotted red lines indicate at which tract locations the groups show significant differences (corrected for multiple comparisons). Tracts are color-coded: Corpus Callosum (CC; green), Superior Longitudinal Fasiculus I (SLF I; light blue), Superior Longitudinal Fasiculus II (SLF II; purple), Superior Longitudinal Fasiculus III (SLF III; dark blue), Inferior Fronto-Occipital Fasiculus (IFOF, pink)

Concerning cortical morphometry, Cortical Thickness (CT) was higher in younger than older adults across large stretches of the cerebral cortex (supplementary material, figure 4). When limiting the areas to the bimanual ROIs, all but the right V5 showed significantly higher CTh in younger than older adults (figure 4).

**Figure 4.**
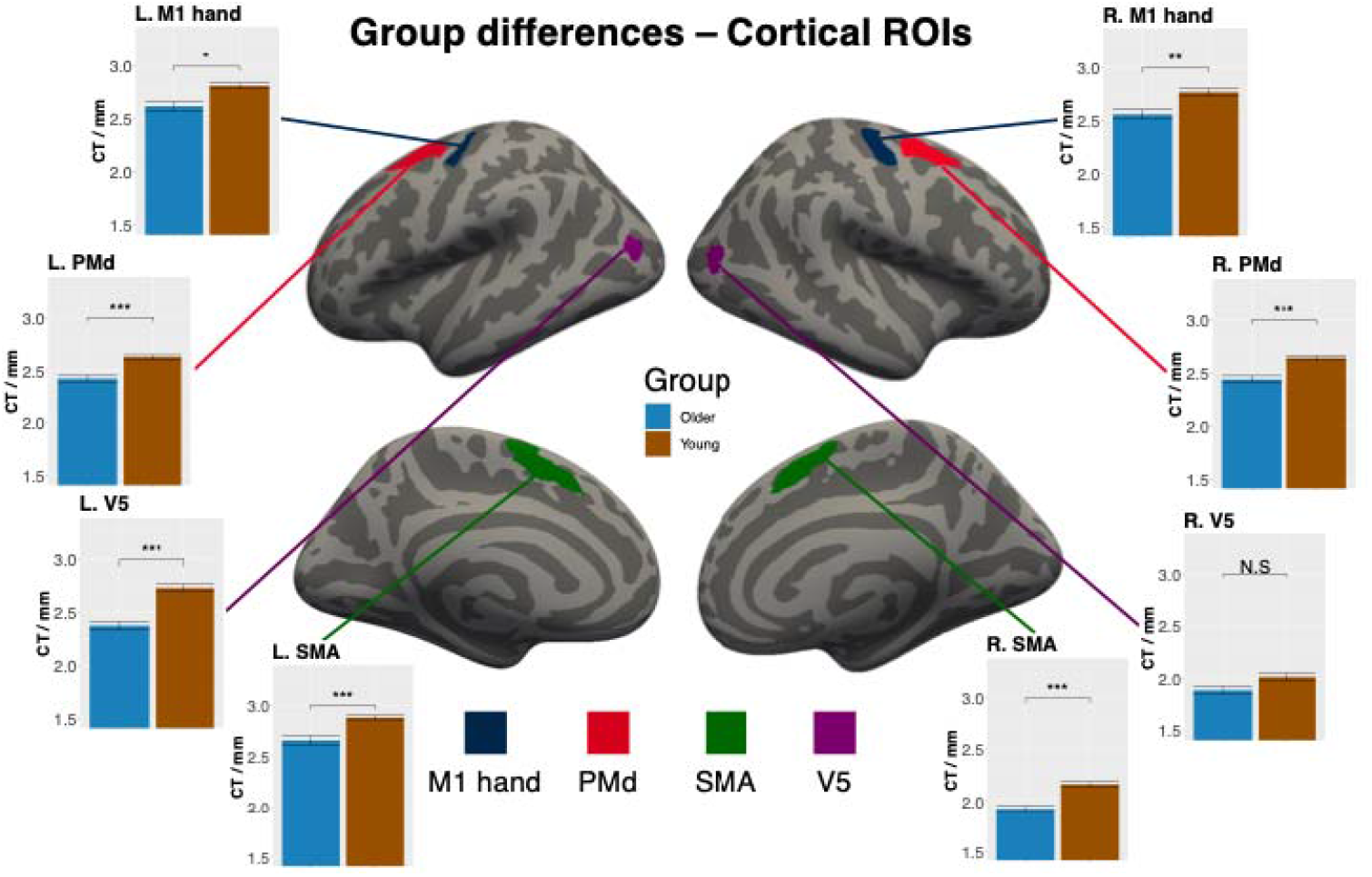
Cortical regions of interest (ROIs) within the bimanual control network. ROIs comprise the hand know of the primary motor cortex (M1 hand; black), dorsal premotor cortex (PMd; red), supplementary motor areas (SMA; green), and middle temporal visual areas (V5; purple). Younger adults (red bars) had a significantly higher cortical thickness (CT) than older adults (blue bars) in all ROIs except for the right V5 area. Significance markers: * p<0.05, ** p<0.01, p<0.001***, Bonferroni corrected between ROIs. L and R denote left and right hemispheres.

### Structural correlates of dynamic bimanual task performance

One of the white tract segments showing age-dependent FA decreases also exhibited a significant correlation with ToT. In the right SLFIII, higher ToT correlated positively with FA in younger but not in older adults (YA: r=0.51; p=0.0198; OA r=0.02; p>0.9) (Figure 5), indicating that higher FA values were related to better performance in the younger adults only. However, correlations were not significantly different between groups, although the p-value approached significance thresholds (z=1.73, p=0.0845). To make sure that the segment identified by group-dependent differences was representative of the structure-performance of the entire tract, we explored the correlation statistics for the entire tract in both younger and older adults. This analysis confirmed that significant positive structure performance relations between the right SLF and ToT could only be detected in younger adults, and r-values were generally higher in younger compared to older adults (YA: mean r-value. 0.193 ± 0.170; OA: mean r-value -0.074 ±0.142; see figure 5 supplementary). To investigate the lateralization of the structure-performance correlations to the right SLF III, we have added the non-significant correlations of the left hemisphere segments of SLF III that showed significant group differences in the supplementary, indicating that the correlations with task performance is specific to the right SLFIII (YA: r=-0.096, p>0.6; OA: r=-0.266, p>0.8) (supplementary figure 6). Correlations between FA and ToT in significant group difference segments are reported in table 2.

**Table 2.** Pearson’s r values and uncorrected p-values for correlations between FA and ToT in all segments showing significant group differences. R-values labeled in **bold** survived multiple comparison correction.

| FA | R. IFOF | R. SLF III | L. SLF III |
| --- | --- | --- | --- |
| Older ToT | $r=0.061$<br>$p=0.391$ | $r=0.024$<br>$p=0.457$ | $r=-0.266$<br>$p=0.891$ |
| Young ToT | $r=-0.092$<br>$p=0.659$ | **$r=0.519^*$**<br>**$p=0.006$** | $r=-0.096$<br>$p=0.664$ |

**Figure 5.**
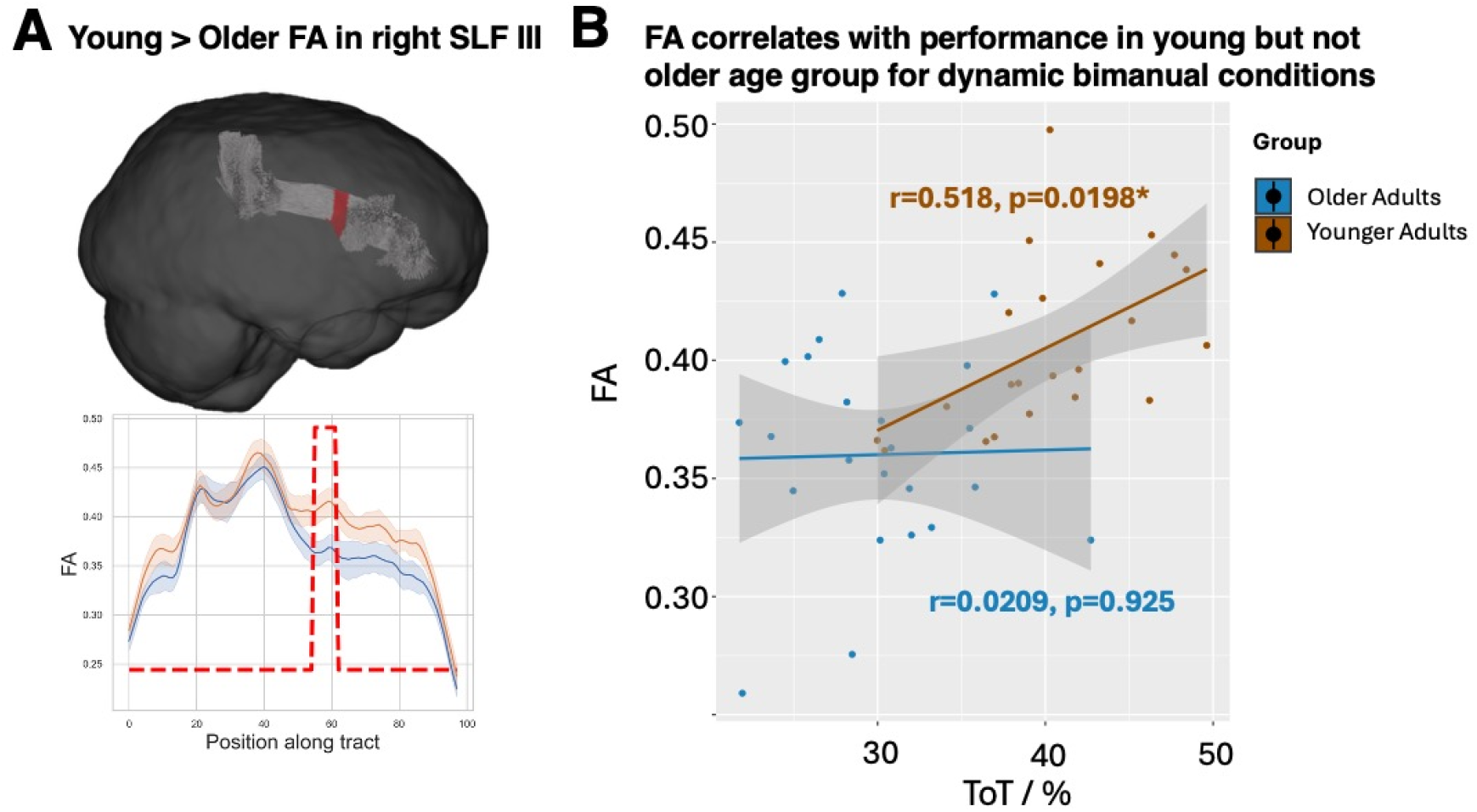
A: Significant group differences in an anterior cluster of the right superior longitudinal fasciculus part III (SLF III). B: Correlations between Fractional Anisotropy (FA) and performance (Time on Target; ToT) within the cluster showing significant age-dependnet effects.

There were no correlations between ToT and MD in any of the significant segments for either of the age groups (p> 0.15). Correlations between MD and ToT in significant group difference segments are reported in supplementary material table 4. For CT, there were also no significant correlations with ToT in any of the ROIs(p> 0.08). Correlations between average CTh in ROIs and ToT are reported in table 4.

**Table 4.** Correlation coefficients and uncorrected p-values for correlations between CTh and ToT in all ROIs. Pearson’s r for all except for L. M1 hand ROI (Spearman’s rho).

| CT | L. M1 hand | L. PMd | L. SMA | L. V5 | R. M1 hand | R. PMd | R. SMA | R.V5 |
| --- | --- | --- | --- | --- | --- | --- | --- | --- |
| Older ToT | $r=0.327$<br>$p=0.074$ | $r=0.068$<br>$p=0.385$ | $r=-0.185$<br>$p=0.211$ | $r=-0.073$<br>$p=0.377$ | $r=0.162$<br>$p=0.242$ | $r=-0.038$<br>$p=0.436$ | $r=0.011$<br>$p=0.482$ | $r=-0.177$<br>$p=0.222$ |
| Young ToT | $r=-0.081$<br>$p=0.364$ | $r=0.197$<br>$p=0.197$ | $r=-0.360$<br>$p=0.0545$ | $r=-0.211$<br>$p=0.179$ | $r=0.392$<br>$p=0.040$ | $r=0.238$<br>$p=0.149$ | $r=0.002$<br>$p=0.496$ | $r=0.206$<br>$p=0.185$ |

### Hemispheric Asymmetries correlate with intermanual symmetry

There was a significant correlation between absolute hand difference in ToT and the absolute hemispheric difference in CTh in the SMA in younger but not in older adults (Figure 6) before correcting for multiple comparisons. Correlations for all cortical ROIs are reported in table 5. However, the difference in correlations between groups was not statistically significant (z=1.21, p=0.227). There were no significant correlations between absolute hand difference and the absolute difference in average FA or MD between the left and right hemispheric homolog tracts (all p values >0.2, see supplementary material).

**Figure 6.**
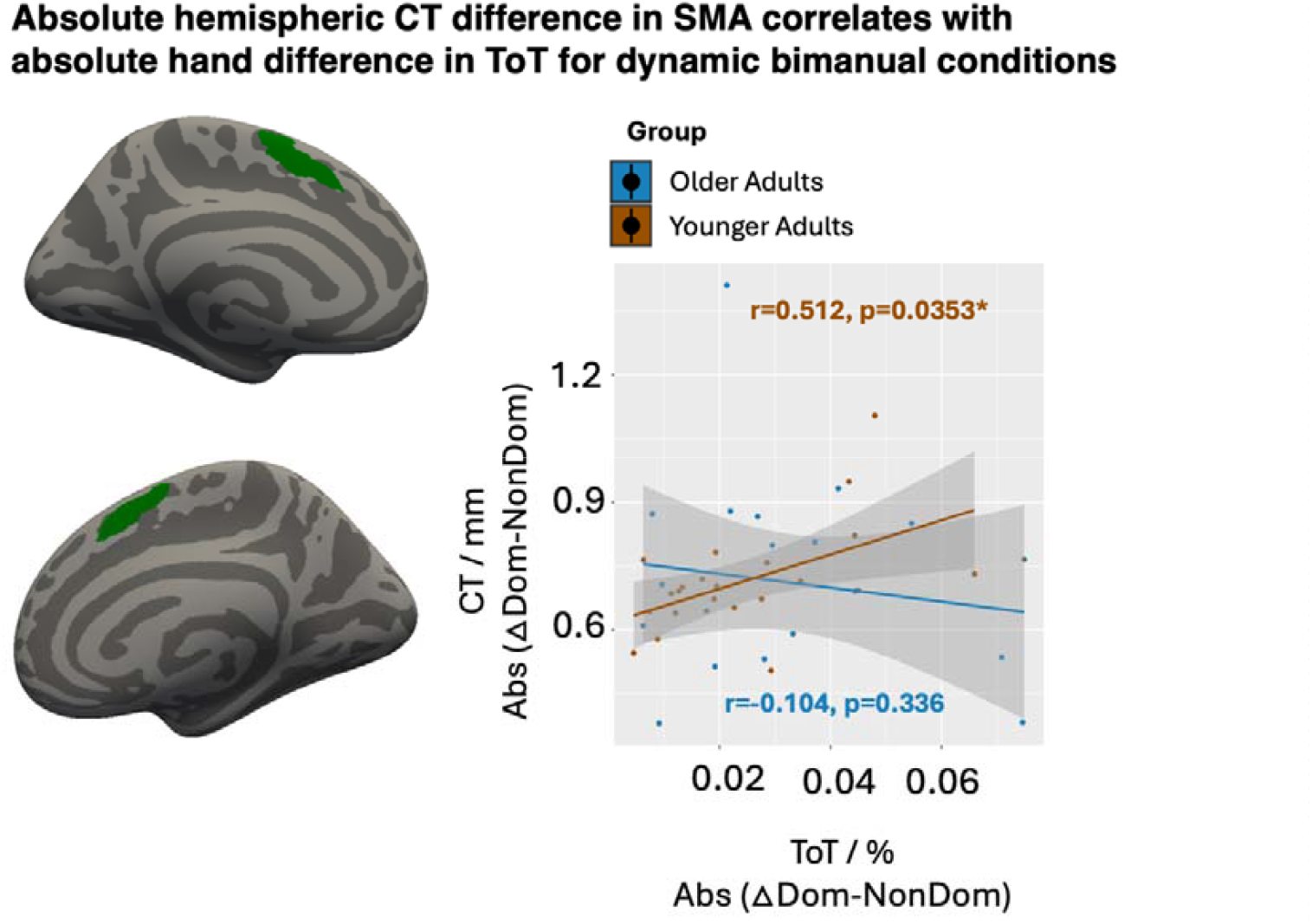
Absolute Time on Target difference between the hands correlates with absolute hemispheric difference in cortical thickness in the SMA in the young but not the old group. On the left, the spatial location of the cluster is visualized. On the right, the correlation between hemispheric symmetry in cortical thickness (CT) (defined as the difference in CT between the dominant (Dom) and non-dominant (NonDom) hemisphere) and hemispheric symmetry in bimanual performance (defined as the difference in Time on Target (Tot) between the Dom and NonDom).

**Table 5.** Correlation values for all hand and hemisphere difference values. Spearman’s rho for all except for V5 ROI (Pearson’s r).

| Abs (ΔDom-NonDom) CTh | M1 hand | PMd | SMA | V5 |
| --- | --- | --- | --- | --- |
| Older abs (ΔDom-NonDom)) ToT | $r=-0.203$<br>$p=0.202$ | $r=-0.200$<br>$p=0.206$ | $r=-0.104$<br>$p=0.336$ | $r=0.376$<br>$p=0.056$ |
| Young abs (ΔDom-NonDom) | $r=-0.047$<br>$p=0.42$ | $r=-0.408$<br>$p=0.033$ | **$r=0.512^*$**<br>**$p=0.009$** | $r=-0.378$<br>$p=0.045$ |

## Discussion

This study investigated how age affects bimanual motor control, white matter microstructure, and cortical thickness within a predefined bimanual motor control network. In addition, it also explored whether structures that showed significant age-related declines in tissue quality correlated more strongly with bimanual performance in older adults compared to younger adults. Our findings corroborate previous literature, showing significantly lower tissue quality in the older adults (Kang & Cauraugh, 2022; Krehbiel et al., 2017; Leiberg et al., 2025, 2025; Roman-Liu & Tokarski, 2021; Rudisch et al., 2020; Vieluf et al., 2015; Yeatman et al., 2014). Younger adults had significantly greater FA values in tracts related to visuomotor processing, specifically in extended sections of the anterior bilateral IFOF and SLF III. These differences in FA were accompanied by significantly lower MD values in similar tract segments. Younger adults also showed significantly higher Cortical Thickness in an extended cluster spanning predominantly frontal, parietal, and temporal regions. Within a bimanual motor network, we found that cortical thickness was significantly lower in the older age group in all but the right V5 area, that is, the bilateral M1 hand, PMd, SMA, and left V5. These structural changes were accompanied by decreased bimanual performance (lower ToT) and inter-hand performance symmetry in older adults.

An exploratory analysis of the relationship between tissue quality and bimanual motor control in regions with significant group differences did not find evidence of stronger structure-performance correlations in older adults. On the contrary, significant correlations were only found in younger adults across behavioral outcomes, and our findings especially highlight the integrity of the right SLF III as functionally relevant in dynamic adaptations in a visually guided bimanual force task. While the correlations were not significantly different in slope when directly comparing the two groups, they may still highlight the functional relevance of intra-hemispheric tracts, particularly the SLF III, for bimanual motor control and offer a basis for future investigation of the functional role of intrahemispheric tracts, leading bimanual research beyond the traditionally emphasized corpus callosum (Fling et al., 2011; Serbruyns et al., 2015).

### Age-dependent changes in the bimanual control network

Our findings show a significant and bilateral age-related decline in FA in the anterior portion of two major association tracts related to visuomotor integration, namely the SLF III and the IFOF. Both are major association tracts related to visuomotor integration, and previous studies have indicated that the volume of these tracts shows positive correlations with good visuomotor control in both younger and older adults (Budisavljevic et al., 2017; Rogojin et al., 2023). The location of the significant age-dependent differences in the anterior segment of the tracts is also aligned with evidence suggesting a general anterior-posterior gradient, with the more frontal white matter showing earlier and more pronounced declines with aging (O’Sullivan et al., 2001; D. Salat et al., 2005). We also found evidence for a widespread, age-related decrease in cortical thickness, with frontal and heteromodal associative regions showing the earliest signs of cortical thinning, which is well aligned with previous literature (McGinnis et al., 2011; D. H. Salat et al., 2004). When focusing on the predefined bilateral motor network, we saw significantly decreased cortical thickness in all ROIs except for the right V5 demonstrating that a significant age-related decrease in cortical thinning is not restricted to frontal and heteromodal regions.

### Age-Dependent Alterations in the Relationship between white matter microstructure and bimanual performance

We were not able to see evidence for stronger structure-function relationships with age within the regions we tested; instead, we found a cluster in the mid-anterior region of the right SLF III that only correlated with performance in the young adults. This is in contrast with most other studies investigating age-dependent alterations in the relationship between while matter microstructure and bimanual coordination, focusing on the relation between callosal integrity and bimanual motor control. These findings suggest that greater white matter integrity between primary and secondary motor regions supports better bimanual motor performance, with age-independent effects (Zivari Adab et al., 2018) as well as age-specific associations favoring older adults (Fling et al., 2011; Serbruyns et al., 2015; Fujiyama et al., 2016). It is noteworthy that most studies highlighting inter-hemispheric connections find that age-specific associations favour older adults, whereas this study, focusing on intra-hemispheric long-range, suggests diminishing structure-function coupling with age. It is worth pointing out that studies outside of the motor control domain have also found evidence for decreased coupling between FA and cognitive performance markers. A study using structural equation modeling on data from individuals from 18–88 years found that the covariance between white matter integrity in intra-hemispheric association tracts and memory performance weakened with age, indicating stronger brain-behavior coupling in younger participants compared to older adults (de Mooij et al., 2018). The authors interpreted this as a sign of age-related dedifferentiation, expressed by a weakening of structure-performance coupling in task-related networks. Dedifferentiation refers to the idea that neural networks become less specialized and more interdependent with age and show reduced functional specificity. It is conceivable that dedifferentiation could impact age-dependent alterations differently in interhemispheric and intrahemispheric tracts, and studies investigating age-related differences in functional connectivity have indicated that while interhemispheric coupling, mediated via the corpus callosum, increases with age, fronto-parietal intrahemispheric coupling decreases (Chen et al., 2019). Taken together these results suggest that intrahemispheric connections may experience different age-dependent effects that are missed if research strongly focuses on the corpus callosum.

The reason for not including the corpus callosum in our correlational investigation is dependent on our experimental approach of only testing in areas where we found age-related changes, and we did not find significant age differences in the truncus of the corpus callosum. This may be explained by the fact that the callosal truncus peaks slightly later (30-40 years) than the criteria for our young age group and decreases most rapidly after 70 y.o. (Hasan et al., 2008; Lebel et al., 2012) and displays a high degree of inter-individual variability in older adults (McLaughlin et al., 2007).

### Anterior cluster in right SLF III correlates with dynamic bimanual task performance in the young group

Higher FA in a mid-anterior cluster in the right SLF III correlated positively with performance in the dynamic bimanual conditions in the young adults. Structurally, the SLF III connects the inferior parietal cortex with the inferior frontal cortex (Janelle et al., 2022). Functionally, its role is lateralized, meaning that the left SLF III is mostly implicated in speech production (Kargar & Jalilian, 2024; Maldonado et al., 2011), while the right SLF III has a crucial role in regulating attention and awareness in visuomanual tasks, such as reaching, grasping, and tool use (Gooijers & Swinnen, 2014; Lazari et al., 2021; Ramayya et al., 2010; Seer et al., 2022; Wang et al., 2016) which fits with our findings highlighting the relevance of the right SLF III in a visually guided pinch-grip task. Studies have shown that the volume of the right SLF II and III correlates with faster visuomotor processing (Budisavljevic et al., 2017). The age effects on this relevance can be nuanced by the observed significant correlation between FA and ToT also when investigating subjects within each group who overlapped in performance, i.e., the older participants who performed better than the lowest performing younger participants and the young participants who performed worse than the highest performing older adult (see supplementary figure 6A). There is still significant lower FA in the segment of SLF III in older subjects in the performance overlap group (see supplementary figure 6B). Judging form the variance of FA in the low-performing older participants, there could be other neural explanations for the very bad performers, possibly detectable in future studies with more participants and with longitudinal designs to study within-subject atrophy and performance decline.

We did not find any significant correlations between CTh in the predefined ROIs of the bimanual motor network and bimanual performance as measured by time on target, and it may be interesting to use structural equation modeling or mediation analysis to measure how the parameters jointly influence behavior. However, the sample sizes required to robustly run such analysis were beyond the sample of this study (Fritz & MacKinnon, 2007; Kline, 2023). Taken together, we observe correlations between bimanual performance and white matter structure in the right SLFIII in the younger but not in the older participants.

### Performance asymmetry between hands correlates with hemispheric asymmetry in cortical thickness of SMA in the younger group

While our main performance measure was a composite score of the left and right hands, due to a strong correlation between ToT measures in both hands, we were also interested in exploring if intermanual symmetry in performance correlated with hemispheric asymmetries in white matter microstructure and cortical thickness. We found that, for younger adults, bimanual symmetry correlated with the absolute difference in cortical thickness in the SMA. This finding agrees with previous fMRI findings from our group in suggesting a crucial role of the SMA in maintaining bimanual symmetry. In the previous work, the blood oxygenation level–dependent (BOLD) signal was recorded in a group of young volunteers during the same task, and the study found that SMA activity, indexed by the BOLD signal, was also predictive of bimanual symmetry during the bimanual pinch-force task (Karabanov, 2023). Together, these two studies show that both structural integrity and functional activity of the SMA are critical for synchronous bimanual control, highlighting the importance of this region in coordinating coordinated motor output across both hands.

### Study limitations and future directions

This study has several limitations. The analyses were exploratory, focused on secondary outcome measures, and based on a relatively small sample size, which may limit both statistical power and generalizability. Accordingly, the findings should be interpreted with caution and warrant replication in larger, confirmatory studies. However, we believe that a key strength of the present work is its emphasis on the relevance of tracts beyond the corpus callosum in associations with (bi)manual motor control. Identifying prospective target tracts like the SLF III is important because studies investigating brain-behavior relationships often limit their analyses to a predefined set of anatomical structures, potentially overlooking critical pathways. This is necessary because brain-wide association studies require very large data sets (N > 100) (Marek et al., 2022). In a resource-intensive technique like neuroimaging, such numbers are often prohibitively expensive and time-consuming, and exploratory work is needed to justify targeted analyses in smaller samples. Moreover, this study was cross-sectional, and more longitudinal studies are needed to make stronger inferences about directionality and potential causality of brain-behavior relationships (Dohm-Hansen et al., 2024).

## Conclusions

In this study, we investigated age-related differences in white matter structure and cortical thickness, focusing on a cortical bimanual motor control network. In regions where age-dependent differences between older and younger adults could be detected, we examined whether aging alters structure–function coupling. We found that a segment in the anterior SLF III was associated with better bimanual performance in younger adults, but we did not find evidence for our initial hypothesis that structure–performance coupling would be increased in older adults. Our findings highlight that fiber tracts beyond the corpus callosum can show significant structure–performance associations. Future studies should investigate whether there are systematic differences between interhemispheric callosal tracts—where increasing associations with age are more consistently reported—and intrahemispheric association tracts, for which our data may offer tentative evidence of decreasing structure–function coupling in older adults.

## Acknowledgments

We thank Keenie Ayla Andersen for assistance with participant recruitment and data collection.

## References

Andersson, J. L. R., Skare, S., & Ashburner, J. (2003). How to correct susceptibility distortions in spin-echo echo-planar images: Application to diffusion tensor imaging. NeuroImage, 20(2), 870– 888. 10.1016/S1053-8119(03)00336-7

Andersson, J. L., & Sotiropoulos, S. N. (2016). An integrated approach to correction for off-resonance effects and subject movement in diffusion MR imaging. Neuroimage, 125, 1063–1078.

B. Fischl, A. Liu, & A. M. Dale. (2001). Automated manifold surgery: Constructing geometrically accurate and topologically correct models of the human cerebral cortex. IEEE Transactions on Medical Imaging, 20(1), 70–80. 10.1109/42.906426

Behler, A., Kassubek, J., & Müller, H.-P. (2021). Age-Related Alterations in DTI Metrics in the Human Brain—Consequences for Age Correction. Frontiers in Aging Neuroscience, Volume 13- 2021. https://www.frontiersin.org/journals/aging-neuroscience/articles/10.3389/fnagi.2021.682109

Boisgontier, M. P., Cheval, B., van Ruitenbeek, P., Cuypers, K., Leunissen, I., Sunaert, S., Meesen, R., Zivari Adab, H., Renaud, O., & Swinnen, S. P. (2018). Cerebellar gray matter explains bimanual coordination performance in children and older adults. Neurobiology of Aging, 65, 109–120. 10.1016/j.neurobiolaging.2018.01.016

Budisavljevic, S., Dell’Acqua, F., Zanatto, D., Begliomini, C., Miotto, D., Motta, R., & Castiello, U. (2017). Asymmetry and structure of the fronto-parietal networks underlie visuomotor processing in humans. Cerebral Cortex, 27(2), 1532–1544.

Chen, Q., Xia, Y., Zhuang, K., Wu, X., Liu, G., & Qiu, J. (2019). Decreased inter-hemispheric interactions but increased intra-hemispheric integration during typical aging. Aging (Albany NY), 11(22), 10100.

Cobia, D., Haut, M. W., Revill, K. P., Rellick, S. L., Nudo, R. J., Wischnewski, M., & Buetefisch, C. M. (2024). Gray matter volume of functionally relevant primary motor cortex is causally related to learning a hand motor task. Cerebral Cortex, 34(5), bhae210.

Cordero-Grande, L., Christiaens, D., Hutter, J., Price, A. N., & Hajnal, J. V. (2019). Complex diffusion-weighted image estimation via matrix recovery under general noise models. NeuroImage, 200, 391–404. 10.1016/j.neuroimage.2019.06.039

Craig, C. L., Marshall, A. L., Sjöström, M., Bauman, A. E., Booth, M. L., Ainsworth, B. E., Pratt, M., Ekelund, U., Yngve, A., & Sallis, J. F. (2003). International physical activity questionnaire: 12-country reliability and validity. Medicine & Science in Sports & Exercise, 35(8), 1381–1395.

Curran, K. M., Emsell, L., & Leemans, A. (2016). Quantitative DTI Measures. In W. Van Hecke, L. Emsell, & S. Sunaert (Eds), Diffusion Tensor Imaging: A Practical Handbook (pp. 65–87). Springer New York. 10.1007/978-1-4939-3118-7_5

Dale, A. M., Fischl, B., & Sereno, M. I. (1999). Cortical Surface-Based Analysis: I. Segmentation and Surface Reconstruction. NeuroImage, 9(2), 179–194. 10.1006/nimg.1998.0395

de Mooij, S. M. M., Henson, R. N. A., Waldorp, L. J., & Kievit, R. A. (2018). Age Differentiation within Gray Matter, White Matter, and between Memory and White Matter in an Adult Life Span Cohort. The Journal of Neuroscience, 38(25), 5826. 10.1523/JNEUROSCI.1627-17.2018

Diedenhofen, B., & Musch, J. (2015). cocor: A Comprehensive Solution for the Statistical Comparison of Correlations. PLOS ONE, 10(4), e0121945. 10.1371/journal.pone.0121945

Dohm-Hansen, S., English, J. A., Lavelle, A., Fitzsimons, C. P., Lucassen, P. J., & Nolan, Y. M. (2024). The’middle-aging’brain. Trends in Neurosciences, 47(4), 259–272.

Dougherty, R. J., Wang, H., Gross, A. L., Schrack, J. A., Agrawal, Y., Davatzikos, C., Cai, Y., Simonsick, E. M., Ferrucci, L., Resnick, S. M., & Tian, Q. (2024). Shared and Distinct Associations of Manual Dexterity and Gross Motor Function With Brain Atrophy. The Journals of Gerontology: Series A, 79(3), glad245. 10.1093/gerona/glad245

Dubbioso, R., Madsen, K. H., Thielscher, A., & Siebner, H. R. (2021). The Myelin Content of the Human Precentral Hand Knob Reflects Interindividual Differences in Manual Motor Control at the Physiological and Behavioral Level. The Journal of Neuroscience, 41(14), 3163. 10.1523/JNEUROSCI.0390-20.2021

Fechner, G. T. (1860). Elemente der psychophysik (Vol. 2). Breitkopf u. Härtel.

Fischl, B., & Dale, A. M. (2000). Measuring the thickness of the human cerebral cortex from magnetic resonance images. Proceedings of the National Academy of Sciences, 97(20), 11050– 11055. 10.1073/pnas.200033797

Fischl, B., Salat, D. H., van der Kouwe, A. J. W., Makris, N., Ségonne, F., Quinn, B. T., & Dale, A. M. (2004). Sequence-independent segmentation of magnetic resonance images. Mathematics in Brain Imaging, 23, S69–S84. 10.1016/j.neuroimage.2004.07.016

Fling, B. W., Walsh, C. M., Bangert, A. S., Reuter-Lorenz, P. A., Welsh, R. C., & Seidler, R. D. (2011). Differential Callosal Contributions to Bimanual Control in Young and Older Adults. Journal of Cognitive Neuroscience, 23(9), 2171–2185. 10.1162/jocn.2010.21600

Fray, P. J., Robbins, T. W., & Sahakian, B. J. (1996). Neuorpsychiatyric applications of CANTAB. International Journal of Geriatric Psychiatry, 11(4).

Fritz, M. S., & MacKinnon, D. P. (2007). Required sample size to detect the mediated effect. Psychological Science, 18(3), 233–239.

Fujiyama, H., Van Soom, J., Rens, G., Gooijers, J., Leunissen, I., Levin, O., & Swinnen, S. P. (2016). Age-Related Changes in Frontal Network Structural and Functional Connectivity in Relation to Bimanual Movement Control. The Journal of Neuroscience, 36(6), 1808. 10.1523/JNEUROSCI.3355-15.2016

Glasser, M. F., Coalson, T. S., Robinson, E. C., Hacker, C. D., Harwell, J., Yacoub, E., Ugurbil, K., Andersson, J., Beckmann, C. F., Jenkinson, M., Smith, S. M., & Van Essen, D. C. (2016). A multi-modal parcellation of human cerebral cortex. Nature, 536(7615), 171–178. 10.1038/nature18933

Gooijers, J., & Swinnen, S. P. (2014). Interactions between brain structure and behavior: The corpus callosum and bimanual coordination. Neuroscience & Biobehavioral Reviews, 43, 1–19.

Hasan, K. M., Kamali, A., Kramer, L. A., Papnicolaou, A. C., Fletcher, J. M., & Ewing-Cobbs, L. (2008). Diffusion tensor quantification of the human midsagittal corpus callosum subdivisions across the lifespan. Brain Research, 1227, 52–67. 10.1016/j.brainres.2008.06.030

Janelle, F., Iorio-Morin, C., D’amour, S., & Fortin, D. (2022). Superior Longitudinal Fasciculus: A Review of the Anatomical Descriptions With Functional Correlates. Frontiers in Neurology, 13. https://www.frontiersin.org/journals/neurology/articles/10.3389/fneur.2022.794618

Jenkinson, M., Bannister, P., Brady, M., & Smith, S. (2002). Improved Optimization for the Robust and Accurate Linear Registration and Motion Correction of Brain Images. NeuroImage, 17(2), 825– 841. 10.1006/nimg.2002.1132

Johansen-Berg, H., Della-Maggiore, V., Behrens, T. E., Smith, S. M., & Paus, T. (2007). Integrity of white matter in the corpus callosum correlates with bimanual co-ordination skills. Neuroimage, 36, T16–T21.

Kang, N., & Cauraugh, J. H. (2022). Bimanual motor impairments in older adults: An updated systematic review and meta-analysis. EXCLI Journal, 21, 1068.

Karabanov, A. N., Chillemi, G., Madsen, K. H., & Siebner, H. R. (2023). Dynamic involvement of premotor and supplementary motor areas in bimanual pinch force control. NeuroImage, 276, 120203. 10.1016/j.neuroimage.2023.120203

Kargar, Y., & Jalilian, M. (2024). Anatomo-functional profile of white matter tracts in relevance to language: A systematic review. Journal of Neurolinguistics, 69, 101175. 10.1016/j.jneuroling.2023.101175

Kline, R. B. (2023). Principles and practice of structural equation modeling. Guilford publications.

Koppelmans, V., Hirsiger, S., Mérillat, S., Jäncke, L., & Seidler, R. D. (2015). Cerebellar gray and white matter volume and their relation with age and manual motor performance in healthy older adults. Human Brain Mapping, 36(6), 2352–2363.

Krehbiel, L. M., Kang, N., & Cauraugh, J. H. (2017). Age-related differences in bimanual movements: A systematic review and meta-analysis. Experimental Gerontology, 98, 199–206.

Lazari, A., Salvan, P., Cottaar, M., Papp, D., van der Werf, O. J., Johnstone, A., Sanders, Z.-B., Sampaio-Baptista, C., Eichert, N., & Miyamoto, K. (2021). Reassessing associations between white matter and behaviour with multimodal microstructural imaging. Cortex, 145, 187–200.

Lebel, C., Gee, M., Camicioli, R., Wieler, M., Martin, W., & Beaulieu, C. (2012). Diffusion tensor imaging of white matter tract evolution over the lifespan. NeuroImage, 60(1), 340–352. 10.1016/j.neuroimage.2011.11.094

Leiberg, K., Blattner, T., Little, B., Mello, V. B. B., de Moraes, F. H. P., Rummel, C., Taylor, P. N., Mota, B., & Wang, Y. (2025). Multiscale cortical morphometry reveals pronounced regional and scale-dependent variations across the lifespan. Cerebral Cortex, 35(6), bhaf154. 10.1093/cercor/bhaf154

Lein Jr, D. H., Alotaibi, M., Almutairi, M., & Singh, H. (2022). Normative reference values and validity for the 30-second chair-stand test in healthy young adults. International Journal of Sports Physical Therapy, 17(5), 907.

Lemaitre, H., Goldman, A. L., Sambataro, F., Verchinski, B. A., Meyer-Lindenberg, A., Weinberger, D. R., & Mattay, V. S. (2012). Normal age-related brain morphometric changes: Nonuniformity across cortical thickness, surface area and gray matter volume? Neurobiology of Aging, 33(3), 617.e1-617.e9. 10.1016/j.neurobiolaging.2010.07.013

Loe, H., Rognmo, Ø., Saltin, B., & Wisløff, U. (2013). Aerobic capacity reference data in 3816 healthy men and women 20–90 years. PloS One, 8(5), e64319.

Madden, D. J., Whiting, W. L., Huettel, S. A., White, L. E., MacFall, J. R., & Provenzale, J. M. (2004). Diffusion tensor imaging of adult age differences in cerebral white matter: Relation to response time. Neuroimage, 21(3), 1174–1181.

Maldonado, I. L., Moritz-Gasser, S., & Duffau, H. (2011). Does the left superior longitudinal fascicle subserve language semantics? A brain electrostimulation study. Brain Structure and Function, 216, 263–274.

Marek, S., Tervo-Clemmens, B., Calabro, F. J., Montez, D. F., Kay, B. P., Hatoum, A. S., Donohue, M. R., Foran, W., Miller, R. L., & Hendrickson, T. J. (2022). Reproducible brain-wide association studies require thousands of individuals. Nature, 603(7902), 654–660.

Matijevic, S., & Ryan, L. (2021). Tract Specificity of Age Effects on Diffusion Tensor Imaging Measures of White Matter Health. Frontiers in Aging Neuroscience, Volume 13-2021. https://www.frontiersin.org/journals/aging-neuroscience/articles/10.3389/fnagi.2021.628865

McGinnis, S. M., Brickhouse, M., Pascual, B., & Dickerson, B. C. (2011). Age-related changes in the thickness of cortical zones in humans. Brain Topography, 24, 279–291.

McLaughlin, N. C. R., Paul, R. H., Grieve, S. M., Williams, L. M., Laidlaw, D., DiCarlo, M., Clark, C. R., Whelihan, W., Cohen, R. A., Whitford, T. J., & Gordon, E. (2007). Diffusion tensor imaging of the corpus callosum: A cross-sectional study across the lifespan. International Journal of Developmental Neuroscience, 25(4), 215–221. 10.1016/j.ijdevneu.2007.03.008

Nasreddine, Z. S., Phillips, N. A., Bédirian, V., Charbonneau, S., Whitehead, V., Collin, I., Cummings, J. L., & Chertkow, H. (2005). The Montreal Cognitive Assessment, MoCA: a brief screening tool for mild cognitive impairment. Journal of the American Geriatrics Society, 53(4), 695–699.

Oldfield, R. C. (1971). The assessment and analysis of handedness: The Edinburgh inventory. Neuropsychologia, 9(1), 97–113.

Oschwald, J., Mérillat, S., Jäncke, L., & Seidler, R. D. (2021). Fractional Anisotropy in Selected, Motor-Related White Matter Tracts and Its Cross-Sectional and Longitudinal Associations With Motor Function in Healthy Older Adults. Frontiers in Human Neuroscience, Volume 15-2021. https://www.frontiersin.org/journals/human-neuroscience/articles/10.3389/fnhum.2021.621263

O’Sullivan, M., Jones, D. K., Summers, P., Morris, R., Williams, S., & Markus, H. (2001). Evidence for cortical “disconnection” as a mechanism of age-related cognitive decline. Neurology, 57(4), 632– 638.

R Core Team. (2021). R: A language and environment for statistical computing (R Foundation for Statistical Computing, Vienna, Austria).

Ramayya, A. G., Glasser, M. F., & Rilling, J. K. (2010). A DTI investigation of neural substrates supporting tool use. Cerebral Cortex, 20(3), 507–516.

Razlighi, Q. R., Oh, H., Habeck, C., O’Shea, D., Gazes, E., Eich, T., Parker, D. B., Lee, S., & Stern, Y. (2017). Dynamic Patterns of Brain Structure–Behavior Correlation Across the Lifespan. Cerebral Cortex, 27(7), 3586–3599. 10.1093/cercor/bhw179

Rogojin, A., Gorbet, D. J., Hawkins, K. M., & Sergio, L. E. (2023). Differences in structural MRI and diffusion tensor imaging underlie visuomotor performance declines in older adults with an increased risk for Alzheimer’s disease. Frontiers in Aging Neuroscience, 14, 1054516.

Roman-Liu, D., & Tokarski, T. (2021). Age-related differences in bimanual coordination performance. International Journal of Occupational Safety and Ergonomics, 27(2), 620–632.

Rudisch, J., Müller, K., Kutz, D. F., Brich, L., Sleimen-Malkoun, R., & Voelcker-Rehage, C. (2020). How Age, Cognitive Function and Gender Affect Bimanual Force Control. Frontiers in Physiology, 11. https://www.frontiersin.org/articles/10.3389/fphys.2020.00245

Salat, D. H., Buckner, R. L., Snyder, A. Z., Greve, D. N., Desikan, R. S. R., Busa, E., Morris, J. C., Dale, A. M., & Fischl, B. (2004). Thinning of the Cerebral Cortex in Aging. Cerebral Cortex, 14(7), 721– 730. 10.1093/cercor/bhh032

Salat, D., Tuch, D., Greve, D., Van Der Kouwe, A., Hevelone, N., Zaleta, A., Rosen, B., Fischl, B., Corkin, S., & Rosas, H. D. (2005). Age-related alterations in white matter microstructure measured by diffusion tensor imaging. Neurobiology of Aging, 26(8), 1215–1227.

Seer, C., Adab, H. Z., Sidlauskaite, J., Dhollander, T., Chalavi, S., Gooijers, J., Sunaert, S., & Swinnen, S. P. (2022). Bridging cognition and action: Executive functioning mediates the relationship between white matter fiber density and complex motor abilities in older adults. Aging (Albany NY), 14(18), 7263.

Serbruyns, L., Gooijers, J., Caeyenberghs, K., Meesen, R. L., Cuypers, K., Sisti, H. M., Leemans, A., & Swinnen, S. P. (2015). Bimanual motor deficits in older adults predicted by diffusion tensor imaging metrics of corpus callosum subregions. Brain Structure and Function, 220(1), 273–290. 10.1007/s00429-013-0654-z

Sexton, C. E., Walhovd, K. B., Storsve, A. B., Tamnes, C. K., Westlye, L. T., Johansen-Berg, H., & Fjell, A. M. (2014). Accelerated Changes in White Matter Microstructure during Aging: A Longitudinal Diffusion Tensor Imaging Study. The Journal of Neuroscience, 34(46), 15425. 10.1523/JNEUROSCI.0203-14.2014

Smith, S. M. (2002). Fast robust automated brain extraction. Human Brain Mapping, 17(3), 143– 155. 10.1002/hbm.10062

Tiffin, J., & Asher, E. J. (1948). The Purdue Pegboard: Norms and studies of reliability and validity. Journal of Applied Psychology, 32(3), 234–247. 10.1037/h0061266

Veraart, J., Fieremans, E., & Novikov, D. S. (2016). Diffusion MRI noise mapping using random matrix theory. Magnetic Resonance in Medicine, 76(5), 1582–1593. 10.1002/mrm.26059

Vieluf, S., Godde, B., Reuter, E.-M., Temprado, J.-J., & Voelcker-Rehage, C. (2015). Practice Effects in Bimanual Force Control: Does Age Matter? Journal of Motor Behavior, 47(1), 57–72. 10.1080/00222895.2014.981499

Wang, X., Pathak, S., Stefaneanu, L., Yeh, F.-C., Li, S., & Fernandez-Miranda, J. C. (2016). Subcomponents and connectivity of the superior longitudinal fasciculus in the human brain. Brain Structure and Function, 221(4), 2075–2092. 10.1007/s00429-015-1028-5

Wasserthal, J., Maier-Hein, K. H., Neher, P. F., Northoff, G., Kubera, K. M., Fritze, S., Harneit, A., Geiger, L. S., Tost, H., Wolf, R. C., & Hirjak, D. (2020). Multiparametric mapping of white matter microstructure in catatonia. Neuropsychopharmacology, 45(10), 1750–1757. 10.1038/s41386-020-0691-2

Wasserthal, J., Neher, P., & Maier-Hein, K. H. (2018). TractSeg—Fast and accurate white matter tract segmentation. NeuroImage, 183, 239–253. 10.1016/j.neuroimage.2018.07.070

Winkler, A. M., Ridgway, G. R., Webster, M. A., Smith, S. M., & Nichols, T. E. (2014). Permutation inference for the general linear model. NeuroImage, 92, 381–397. 10.1016/j.neuroimage.2014.01.060

Wulff-Abramsson, A., Zvornik, A., Andersen, K., Yang, Y., Novén, M., Lundbye-Jensen, J., Tomasevic, L., & Karabanov, A. N. (2025). Event-related theta synchronization over sensorimotor areas differs between younger and older adults and is related to bimanual motor control. NeuroImage, 121032. 10.1016/j.neuroimage.2025.121032

Yeatman, J. D., Dougherty, R. F., Myall, N. J., Wandell, B. A., & Feldman, H. M. (2012). Tract Profiles of White Matter Properties: Automating Fiber-Tract Quantification. PLOS ONE, 7(11), e49790. 10.1371/journal.pone.0049790

Yeatman, J. D., Wandell, B. A., & Mezer, A. A. (2014). Lifespan maturation and degeneration of human brain white matter. Nature Communications, 5(1), 4932.

Zivari Adab, H., Chalavi, S., Monteiro, T. S., Gooijers, J., Dhollander, T., Mantini, D., & Swinnen, S. P. (2020). Fiber-specific variations in anterior transcallosal white matter structure contribute to age-related differences in motor performance. NeuroImage, 209, 116530. 10.1016/j.neuroimage.2020.116530

Zvornik, A., Andersen, K. A., Petersen, A. D., Novén, M., Siebner, H. R., Lundbye-Jensen, J., & Karabanov, A. N. (2024). Older and younger adults differ in time course of skill acquisition but not in overall improvement in a bimanual visuomotor tracking task. Frontiers in Aging Neuroscience, 16. 10.3389/fnagi.2024.1373252

